# An entropic barriers diffusion theory of decision-making in multiple alternative tasks

**DOI:** 10.1101/245308

**Authors:** Diego Fernandez Slezak, Mariano Sigman, Guillermo A. Cecchi

## Abstract

We present a theory of decision-making in the presence of multiple choices that departs from traditional approaches by explicitly incorporating *entropic barriers* in a stochastic search process. We analyze response time data from an on-line repository of 15 million blitz chess games, and show that our model fits not just the mean and variance, but the entire response time distribution (over several response-time orders of magnitude) at every stage of the game. We apply the model to show that (a) higher cognitive expertise corresponds to the exploration of more complex solution spaces, and (b) reaction times of users at an on-line buying website can be similarly explained. Our model can be seen as a synergy between diffusion models used to model simple two-choice decision-making and planning agents in complex problem solving.

## 1 Introduction

Decision-making has been studied in great detail relying on binary choices by the Two-Alternative Forced-Choice paradigm (2AFC). In 2AFC tasks, choices are made between two alternatives with limited information while speed and accuracy are registered. In addition, for simplicity, in the vast majority of the experiments, the decision variable is a single scalar (for example the luminosity of patches, the number of dots in a comparison task or the pitch in an auditory discrimination task).

This paradigm has a great benefit for computational and theoretical understanding of decision making. It can be fully expressed in a set of equations which have analytic solution [1]. Also, the functional dependence of behavioral observables such as response times, error rates or confidence can be described in detail by low dimensional models (e.g. [1–5]). While models differ, many of them rely on a stochastic search process, in which the accumulation of the evidence is integrated over time, and whose crossing of a boundary represents the event of *reaching a conclusion* or making a decision [6–8].

The Drift Diffusion Model (DDM) [2, 9, 10] has been shown to be, under some conditions, optimal mechanism model for 2AFC decision-making [7, 11]. The discrete analogue of DDM consists of a random walk on an 1-D interval, with one extreme as origin and the other as the decision boundary. Several variants of this model have been proposed depending on how the threshold is set, whether the integrated signal decays in time, and whether the two choices are represented by two competing (and possibly interacting) signals. More recently, the Ising Decision Maker has been presented as a new formal model for 2AFC, showing increased performance compared to Ratcliff diffusion model [12].

Over many years, this research program has shown with exquisite detail how humans and other animals reach decisions with a small number of options. However, most real-life decisions are made of a large (often infinite) number of choices, relying on heuristics and a (relatively shallow) search process in decision trees with complex geometries.

Beyond some simple scenarios where classic diffusion models can be extended to more dimensions, this class of models can hardly adequately describe decision-making in multiple choices [13, 14]. For instance, Usher and McClelland introduce the leaky, competing accumulator model extending this framework to multiple-decision tasks [5]. This model proposes several leaky integrators of signals – based on Ornstein-Uhlenbeck equations – which compete and inhibit each other until a decision is made by reaching a threshold. This model has a greater number of parameters (increasing the degrees of freedom) and show similar patterns of RT curves as DDM, which lead to (small) systematic gains (in terms of quadratic error in fitting) compared to DDM results.

This tree-search approach has been studied broadly in Cognitive Psychology. Lee and colleagues have successfully used DDM in sequential 2AFC tasks where context conditions are changed during time, and showed how diffusion models with non-homogeneous information focus on search for evidence and explain adaptation on search termination [15]. Another example are Multi-modal Processing tree models is a framework for developing and testing quantitative theories based on observations in categorical data [16]. It provides a data-analysis tool capable of disentangling and measuring separate contributions of different cognitive processes, measuring latent processes that are confounded in observable data. However, as MPT models are explanatory they require a detailed description of cognitive processes behind the behaviour under study, which may be impossible to propose when investigating high-level phenomena.

In the artificial intelligence literature, techniques from operations research have been used to attack problems of choosing actions by planning agents in partially observable stochastic domains [17–19]. These methods build a policy tree – where actions are selected to optimize reward – and resemble the monte-carlo searches used to model the dynamics of decision-making.

The objective of the present work can be viewed as an effort to bring together diffusion models in binary decisions and planning agents in complex problem solving, two very influential, but largely disconnected literatures of decision-making. We theorize that the defining characteristic of a generic decision process – in humans and computer algorithms – is the presence of *entropia barriers* [20], i.e. paths that are diffusively explored and usually lead to dead ends or sub-optimal solutions.

We investigate here whether the distribution of reaction times (RT) in multiple-alternative decision-making can be modeled by a diffusion process on a space with topological traps, as we intend to capture the essence of a stochastic search process involving the exploration of “dead ends” and the concomitant back-tracking. In other words, we hypothesize that a random walk in a 2-D grid with obstacles may represent the decision process trajectories as tree search algorithms. Each position in the grid represent different board states, radius of bound reflects search depth, obstacle density reflects amount of pruning and time per step represents processing speed. We emphasize that while the model aims to integrate in its simplest form the notion of diffusion to a bound with the idea of exploring branches in a tree, it is not intended to be an exact correspondence, as the topology of all the decisions problems in the search tree cannot be embedded in a one to one fashion in the 2 dimensional grid.

A traditional limitation of behavioral and cognitive modeling has been the mismatch between the complexity of the models and the availability of experimental data to validate them. The advent of big data, however, has turned this difficulty on its head [21]. To examine our hypothesis, therefore, we capitalized on a vast corpus of decision-making obtained from on-line chess servers [22]. This database has several virtues: (1) in any given move, players have to opt among a large number of options [23], (2) options grow exponentially (three steps down on decision tree typically results in more than a billion alternatives) and hence the search process becomes rapidly intractable without heuristics, (3) as in real-life, in chess choices have to be made with a finite time budget (4) the quality of the decision-maker is particularly easy to calibrate in chess (players have an ELO which indicates the quality of their decisions), and (5) it contains detailed information about more than 1 billion decisions, a volume that would be unthinkable to reproduce in a classical laboratory setting.

We show that this simple theoretical construct, a natural extension of diffusion processes with the inclusion of obstacles (or entropic barriers) provides a remarkably accurate description of the data that was not captured with previous models. We also show that this obstacles model may characterize individual RT distributions and predict similarity with other players based on their time-to-move distributions. Moreover, we test the model in a complete different scenario of an online buying website where, suggesting that this model may describe a broad class of decision-making processes ranging from simple binary decision-making to complex decision in problem solving with heuristics not only in chess playing.

## 2 Methods

### 2.1 Drift-Diffusion Model

The Drift Diffusion Model (DDM) is the continuous analogue of a random walk with a direction bias, and is considered the optimal model for simple two-choice decision processes [11]. The model assumes that decision is made by the integration of evidence in a noisy process over time. Our implementation consists of a particle in a 2-D mesh walking to a threshold. Steps may be in any of four directions (up, down, left, right) equally distributed. Bias is inserted by accepting the step following the conditions:

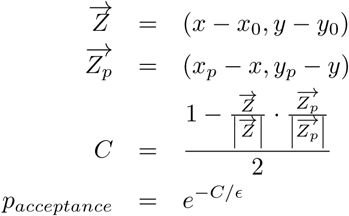

where 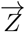 is the current position, 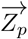 is the intended next position, *C* is the cost of putative move, *p* the probability of accepting the move and *ε* the noise in the acceptance decision. Thus, the parameters of the simulation are the radius of the boundary (in units of the grid step) *R*, the decision noise ∊ and the time step Δ*t*.

Several alternatives to DDM model appear in the literature, both for 2AFC and multiple-choices extensions (e.g. [5, 12]). These models show similar RT patterns and increase the number of free parameters, which permit (small but systematic) gains in fitting RT distributions. However, in our complex chess scenario, patterns of RT distribution differ significantly from these classic models. Thus, for sake of clarity we will compare our obstacles model to DDM.

### 2.2 Obstacles Model

A one-dimensional random walk [24], representing a blind decision-making process, initiates its exploration at a fixed position. The walker ends its exploration when it reaches a threshold value, after which the process may be re-initiated. The time to reach this threshold is the first-passage time (FPT), and characterizes stochastic models of RT.

In continuous time and space, the dynamics are expressed as:

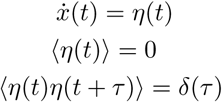

The transition probability *P*(*x*|*x', t)* of finding the walker in *x* at time *t* given that it was in *x*′ at time 0 satisfies the Fokker-Planck equation

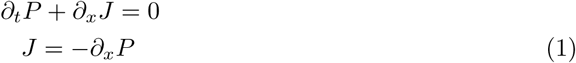

on which the initial and boundary conditions are easily expressed: assuming the origin of the walker at *x*_0_, the initial condition implies *P(x|x*_0_, 0) = *δ(x − x*_0_), a crossing threshold at *x_T_* is equivalent to an absorbing boundary, *P(x_T_|x*_0_*, t)* = 0, whereas a reflecting boundary at *x_R_* corresponds to *J*(*x_R_* |*x*_0_, *t*) = 0. The probability distribution for the FPT is

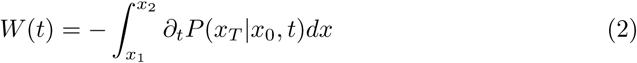

where *x*_1_ and *x*_2_ are the boundaries (one is the absorbing, the other may be absorbing, reflecting or located at ∞), and *x*_1_ < *x*_0_ < *x*_2_. The analytic solution to this problem is known, and the limiting cases of reflecting boundary at finite and at infinite distance result in exponential and power-law tail distributions, respectively. The extension to higher-dimensional Euclidean geometries of the analytic results demonstrate a similar behavior for the tail distributions ( [25], see also Suppl. Mat.).

These kind of models have been successfully used to model 2AFC and other tasks. However, we show that these models fail to represent the full distribution in more complex scenarios such as RT in chess and online buying. We propose that including entropic barriers in a 2D space would add the neccesary complexity to the model so that it would replicate the full distribution. Opposely to FPT models, the 2D space with entropic barriers has no analytic solution. Thus, we simulate the model by a random walk, discretizing the 2D space into a discrete mesh where some of the nodes are considered entropic barriers and the random walk is not able to pass through.

In the obstacles model, we use a square grid with a circular absorbing boundary, and reflecting nodes scattered randomly with a given density. We simulate trajectories in the 2-dimensional plane, since this corresponds to the minimum number of dimensions in which obstacles do not necessarily disconnect space. The process moves randomly and without bias in one of the four possible grid directions at each step. The parameters of the simulation are the radius of the boundary (in units of the grid step) *R*, the smoothness of the space *ρ* (i.e. the number of obstacles in the grid), and the time step Δ*t*. The update rule is then for position 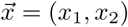 at time *t* is:

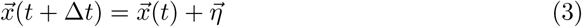

where the random drift 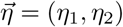 is defined formally by *η_i_* = ±1, 〈*η_i_*〉 = 0, 〈*η_i_η_j_*〉= *δ_ij_.*

The smoothness is related to the probability that a grid point is an obstacle, 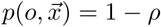, 0 ≤ *ρ* ≤ 1, so that *ρ* = 1 is a pure Euclidean space. The obstacles are defined by the constraint on the probability current, 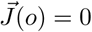. Figure 1 shows a walker that starts in the center and wanders through the labyrinth with obstacles until it finds a passage point.

**Fig 1.**
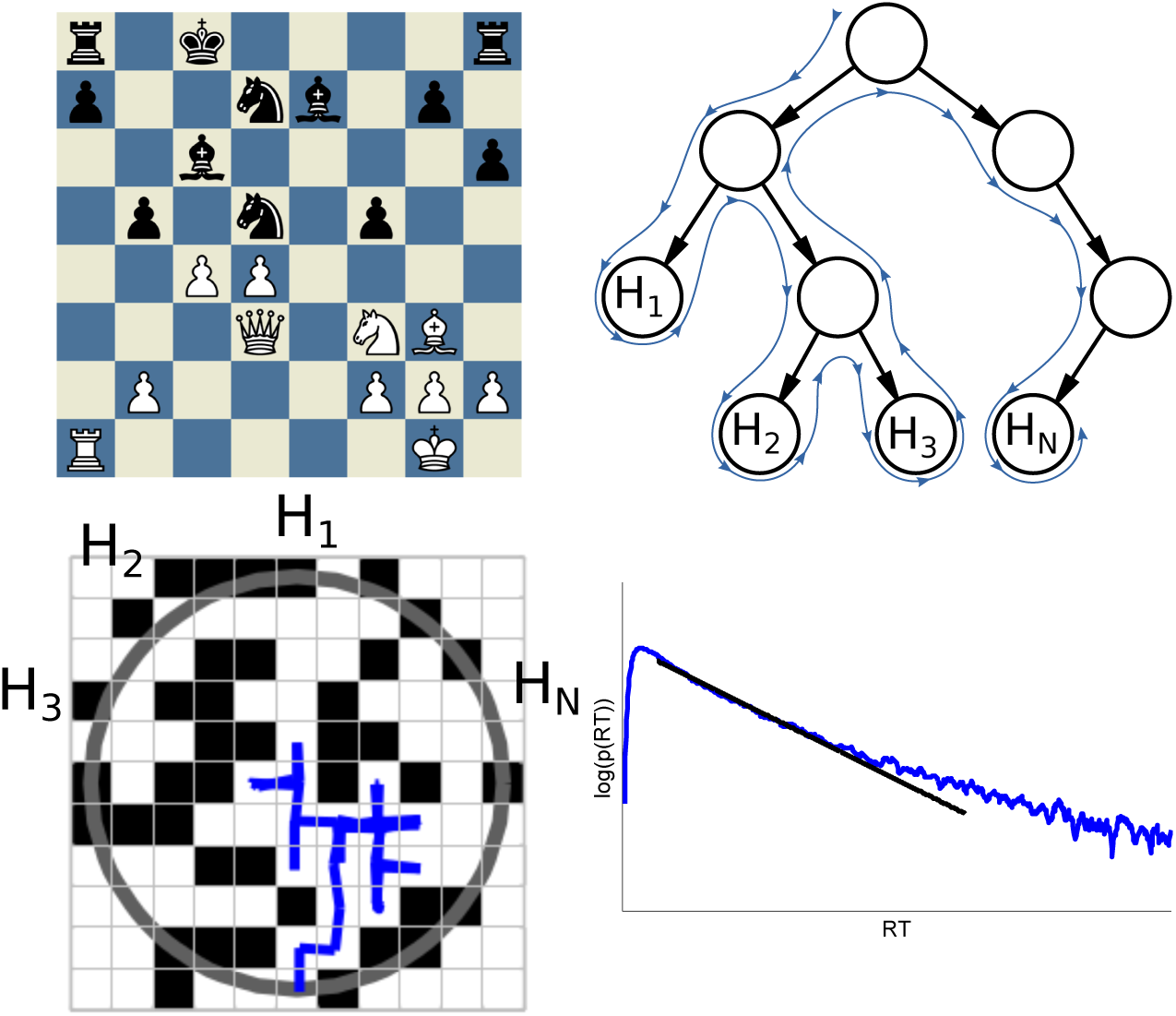
a) Chess board example of the game between Gary Kasparov and Deep Blue on May 11th, 1997 – the winning game of machines against humans. b) Decision tree for current position and hypothetical walk through it until decision is made. c) Particle moving in a 2-D mesh with obstacles resembling the tree search heuristic. d) RT distribution with super-exponential tail.

### 2.3 Fitting RT Distributions

Simulations of models were implemented in Python. Each execution of a model with fixed parameters returns a number of steps taken to reach the threshold which represents a response time of a single decision.

Parameters fitting of both models (DDM and obstacles model) was performed by exhaustive search in discrete parameters (distance) and Nead-Melder method [26] for non-linear optimization over the continuously parameter space (step size and number of obstacles). For each parameter combination, we executed 75.000 simulations, i.e. 75.000 simulated decisions. In the case of the DDM, the parameter space range was: *R* ∈ [1, 10] in grid-steps units, *∊* ∈ [0, 0.25] and Δ*t* ∈ [10, 150] in milliseconds. On the other hand, the obstacles-model parameter space range was: *R* ∈ [1, 10] in grid-steps units, *ρ* ∈ [0.2, 0.65] and *Δt* G [10, 150] in milliseconds.

To fit the human RT distribution, for each parameter combination we calculated the Jensen-Shannon divergence (JSD) [27] between the human *(hRT*) and the model *(mRT*) distributions. Thus, the optimization method consists on the parameters lookup in the simulated-decisions distributions which minimizes:

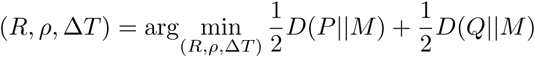

where *M* = 1/2(*P* + *Q*), *D(P*||*M*) the Kullback-Liebler divergence [28], *P* and *Q* the distributions to compare.

To compare the relative goodness of fit between our model and DDM, we also use JSD instead of the usual AIC, following [29].

## 3 Results

We compiled RT statistics of more than 15 millions of chess games (i.e. more than 1 billion decisions) played between 2006 and 2013 by human players with a total time budget of 300 seconds. These games were obtained from the Free Internet Chess Server^1^, a free ICS-compatible server for playing chess games through Internet, with more than 300.000 registered users. This massive dataset also includes the user rating at each game – a dynamic variable which is updated after each game played according to the Glicko method [30] – that indicates the chess skills strength of the player.

We computed the 2-D histogram of players RTs as a function of the seconds of games played – or similarly as a function of remaining time (see Fig. 2a). As observed in our previous work [22], decision times are shorter during the first and last stages of the game, caused by the standardized openings or time constraints of the endgame (see Fig. 2a). Here, we consider only the middle stage of the game, defined as all moves performed when fraction of remaining time is between 0.1 and 0.9, i.e. total time remaining is between 270 seconds and 30 seconds. All the RT distributions in this middle stage present a very similar shape: starting from a few almost immediate moves (less than half-a-second RT), a rapid growth to the mode RT (mode value depends on total time and remaining time), with a super-exponential decaying tail.

**Fig 2.**
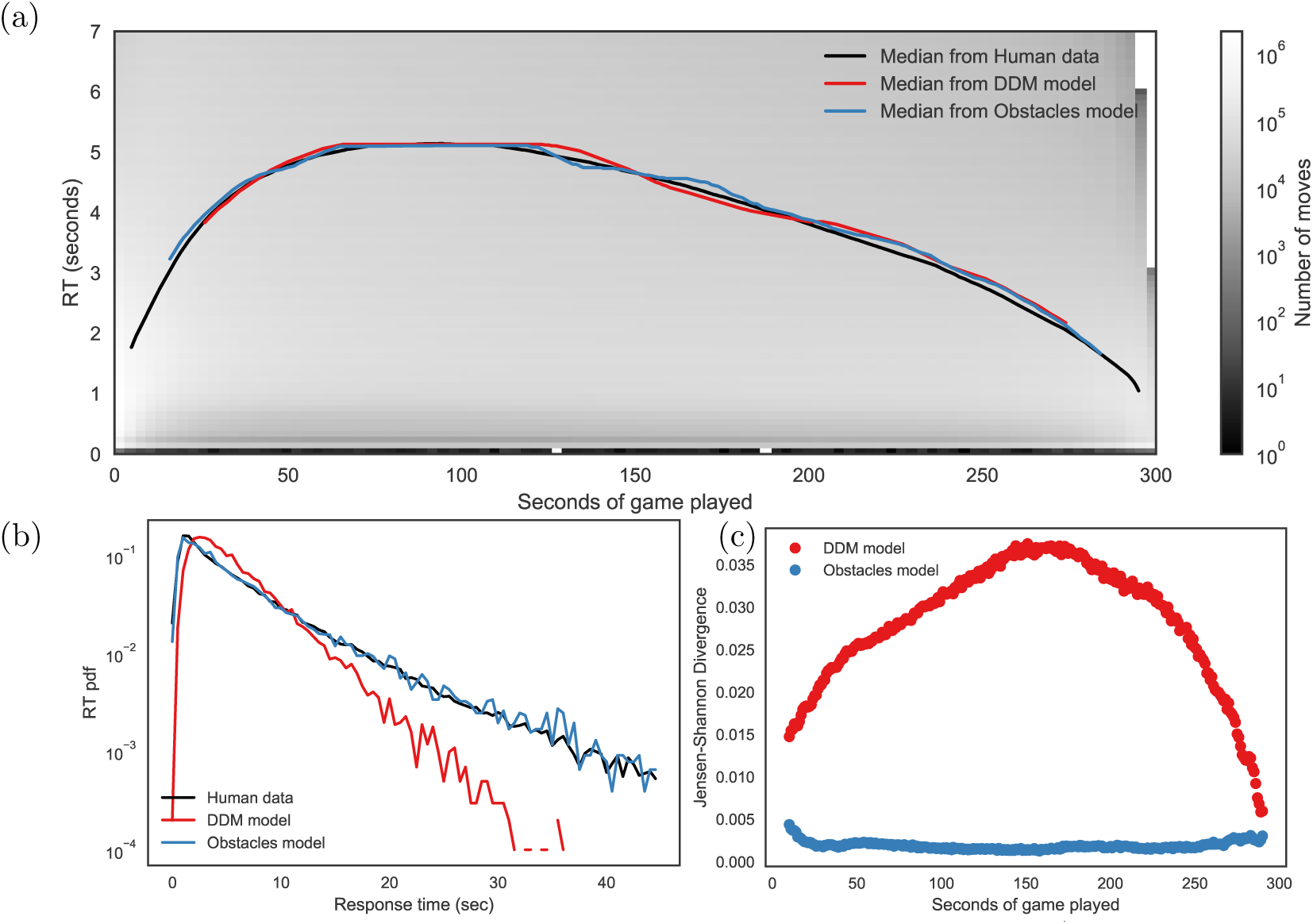
Response Time statistics and fit for human players. a) 2-D histogram (in log scale) of RT of moves as a function of the seconds of game played. Lines show median RT for human statistics (black line) and best fit with no entropic barriers (red line) and obstacles model (blue line). No significant difference between distributions at every instant in both models, Kolmogorov-Smirnoff Test, *KS_statistic_* < 10^−4^, *p* > 0.99). b) RT distribution at 50% of the game (black trace). Model with obstacles (blue line) fits with extremely high precision in both short and long time scales (Kolmogorov-Smirnoff statistics *D_n_* = 0.014, *p* > 0.95). The red line is the best fit for the DDM (Kolmogorov-Smirnoff statistics, *D_n_* = 0.754, *p* < 10^−6^)). c) Goodness of fit indicated by Jensen-Shannon divergence. Model without obstacles (red dots) shows significantly higher values than proposed obstacles model (blue dots) at every instant of the game.

### 3.1 RT distribution models

We fitted parameters for both models (DDM and obstacles model) for the RT distribution obtained at each instant of the game. That is, given a remaining time, we estimated the best parameters which make the model fit more accurately the RT distribution at that instant of the game. The remaining time was divided into 0.1 seconds bins, i.e. 3000 bins in 300 seconds of total time available per game. Once this parameter fitting was performed for each instant of the game, we calculated the estimated median of the RT distribution (Fig 2b). At every instant of the game, both models – with obstacles (blue line) and classic DDM (red line) – show an extremely precise median (KS Test, no significant difference between distributions at every instant).

However, if we compare the goodness-of-fit of the complete distribution we find that obstacles model fits significantly better than classic DDM. As an example of this, we show the parameter fitting of RT distribution at 50% of game using a single DDM model with only 3 free parameters: *R* the threshold distance, ∊ noise in the acceptance decision, and Δ*T* the time per step (see Methods section for fitting details). Standard diffusion models (in particular, DDM without obstacles) produce distributions of FPT with exponential decay or power-law regardless of the dimensionality ( [31], see also Suppl. Mat.). As expected, this fit is inconsistent with our observations of RT distributions which display a similar initial distribution but a super-exponential tail. We observe the best fit for RT distribution at the middle of the game using a single DDM (∊ = 0.2, *R* =8 and Δ*T* = 60ms, see Fig. 2c, red line) which is numerically very distant to the data (Kolmogorov-Smirnoff statistics, *D_n_* = 0.754, *p* < 10^−6^) and, moreover, shows a qualitative different behavior with a tail that decays more rapidly. Instead, fitting with the 2-D diffusion model with obstacles (see Fig. 2c, blue line) results in a very accurate description of the data (Kolmogorov-Smirnoff statistics *D_n_* = 0.014, *ρ* > 0.95). For this representative fit, *ρ* = 0.45 (45% of grid points are obstacles), with a radius of *R* = 4 and Δ*T* = 90ms.

We repeated this procedure for each instant of the game and calculated the goodness of fit of both models, estimated by the Jensen-Shannon divergence. We observe that model with obstacles outperforms classic DDM (Fig 2d). Model without obstacles shows values of JS divergence much higher than obstacles models in every instant of the game.

The model is reminiscent of “comb” geometries, consisting of a main backbone with the origin and threshold in each extreme, and side branches where the diffusing particle may get trapped. This configuration is more restrictive in the topology of search process, but a comparative advantage is that analytic solutions for this problem have been developed. However, the best comb solution does not fully capture the observed RT distributions (see Supp. Mat. for details).

The results above show that diffusion with obstacles model provides an accurate description of RT distributions obtained from a vast corpus of human decisions. However these distributions aggregated decisions from many different players. Hence a possible and alternative explanation is that the non-exponential tails we observed in the data resulted from an addition of different exponentials (assuming that players may have different characteristic decision times). To test this hypothesis, we selected individual players with more than 20.000 games each (17 players in total) and performed the model fitting considering entropic barriers for each individual player. The model could fit very accurately the distribution of RTs of each individual player, reflected in small KS-statistics and p-values close to 1, which indicate that the model and the data are not distinguishable (KS test, *p* > 0.99, *D_statistic_* < 10^−8^ in all cases; see fits in Supp. Mat.). This shows that the inability to fit the distribution of all decisions was not a matter of aggregating different individuals: instead the model with entropic barriers provides a very accurate fit of the RT distribution of individual players.

### 3.2 Model performance and comparison

To compare models, we implemented a cross-validation scheme. For each player with high number of games, we selected the RT distribution at 50% of the game and splited into 80%-20% sets for training both models and testing the fits, respectively. Using the first set, we fitted both models parameters and evaluated in the test set, obtaining 2 JSD measures, one for each model. We repeated this procedure 1000 times and calculated the average JSD values for each model. Obstacles model showed better performance than DDM model in average JSD for all players (see Supp. materials for details).

Then, we evaluated the prediction capabilities of the model. We proposed that players who show similar RT distributions (in terms of JSD values), should obtain similar JSD values for the fitted distributions of the model. Thus, given both RT distributions of a particular player (the real data, and the fitted model), sorting other players based on similarity to their RT distribution must correlate. We fitted each player individually and calculated JSD values of real data and the fitted model to every other player. We found that correlation between sortings is significantly better using the Obstacles model (Spearman rank-correlation *r* > 0.99, *p* < 10^−10^ in all cases) than DDM (Spearman rank-correlation *r* ≈ 0.6, *p* < 10^−2^ in most cases, with some iterations not significant).

Our next aim is to test the Obstacles model, examining *specificity,* i.e. whether it can be used to obtain useful discriminations and *generality,* i.e. whether the model is valid across different contexts.

### 3.3 Specificity

To test specificity we investigated whether it can identify meaningful differences in the decision process between good and bad players. A comparative advantage of studying decision-making in chess is that the quality (i.e. rating) of players can be precisely determined, representing the player’s strength in a very confident manner. The rating of a player is determined based on their past results with other players, and updated after each game played. Players receive rating points in proportion to the difference in their strength. A strong player would increase a very small amount of rating points when winning a game versus a weak player. On the contrary, a weak player would win rating points even in the case of a draw playing to a stronger opponent [30].

Decision-time distributions of strong and weak players are comparable, although stronger players make slightly slower moves during middle-game and faster during the opening and the end-game [22]. We reasoned that this subtle differences in RT distributions between strong and weak players may rely on different decision processes. Specifically, in line with notions of expertise [32] we hypothesized that good players (expert decision-makers) discard a larger number of states which through heuristic which in our model accounts to having a larger number of obstacles (i.e *ρ_goodplayers_* > *ρ_badplayers_*) [33] which is compensated by a faster navigation of the decision-tree (i.e Δ*T_goodplayers_* < Δ*T_badplayers_*). As for the radius (the equivalent to the depth of search), different chess theories differ on whether search depth increases with quality or remains constant.

To examine these hypotheses we splited the population into quintiles groups (i.e. each group has the same number of played games) and analyze distinctively the parameters of the model. We performed the parameter fit to data corresponding to each one of the five groups. The correlation between each parameter (obstacles rate (*ρ*), time per step (Δ*T*) and threshold distance (*R*)) and player ratings are presented in Fig. 3. With higher ratings, players show higher obstacles ratio (Fig. 3a, Pearson correlation *r* = 0.89, *p* < 0, 04) and shorter distances (Fig. 3c, Pearson correlation *r* = −0.97, *p <* 0, 006). Time per step showed no correlation with player ratings.

**Fig 3.**
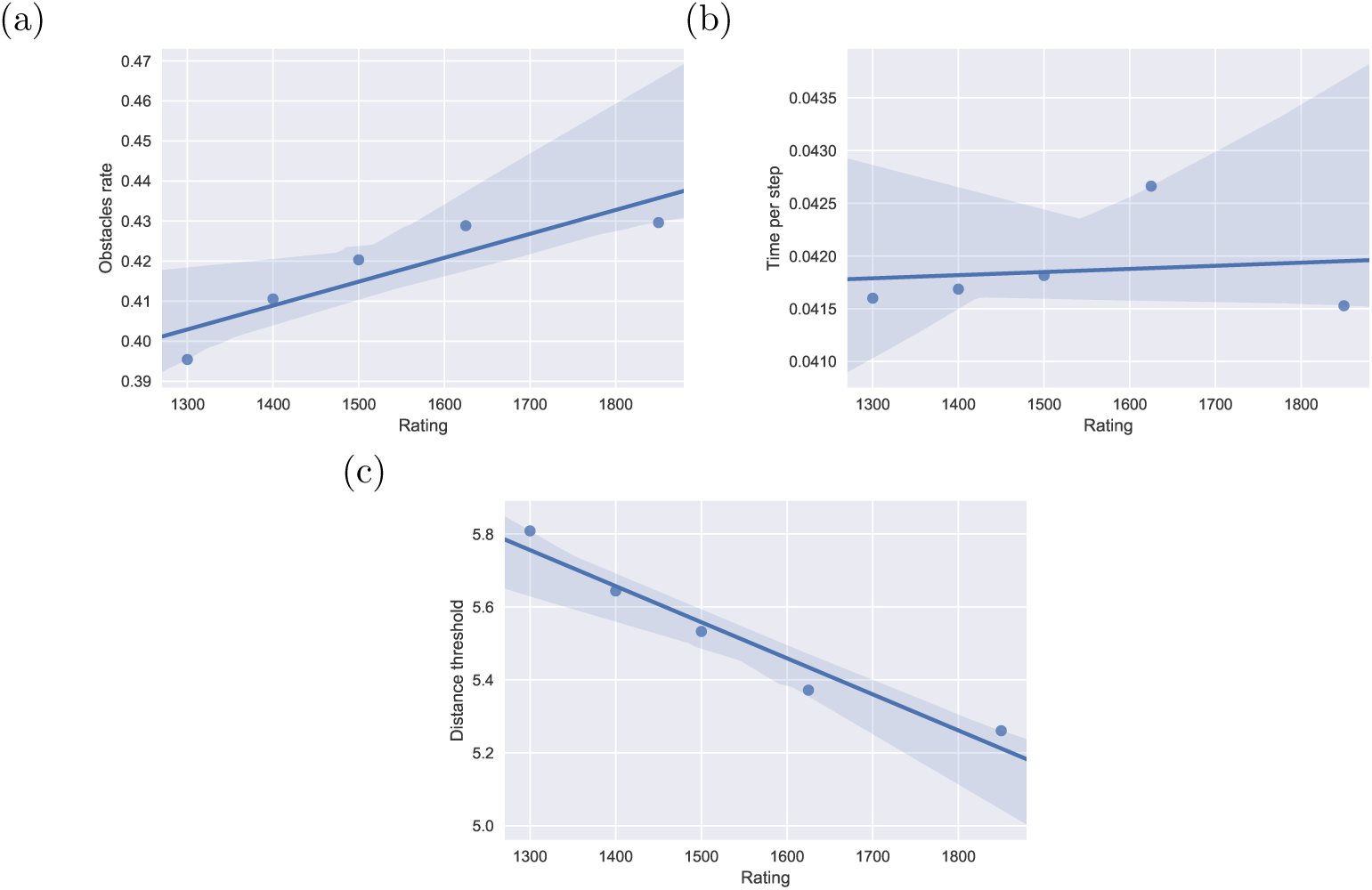
Specificity. Correlation between parameters of the model and player ratings. a) Density of obstacles (a proxy for the complexity of the space of solutions explored), b) time per iteration and c) distance to threshold as a function of time in the game. Correlations show significant p-values for density of obstacles (*p <* 0.006) and distances (*p* < 0.04). Strongest players (first quintile) explore a more complex decision space (more obstacles, KS Test *p <* 10^−17^)) compared to weakest players (fifth quintile), with similar time per step (KS Test, *p* = 0.20), leading to shorter distances (KS Test, *p <* 10^−7^).

Obstacle density show higher values for strongest players compared with weakest players (KS Test, *p <* 10^−17^), showing more complex grids to transit during the random walk. On the other hand, time per step show similar patterns between both classes of players (KS Test, *p* = 0.20), suggesting that this parameter may be representing a canonical aspect of decision making which does not differ among player classes. As strong players show more complex scenarios with similar time per step, distances to threshold show smaller values in strong players than in weak players (KS Test, *p <* 10^−7^). These results suggest that all players spend the same time for checking each step, but strong players explore more difficult decision spaces (more obstacles), with shorter distances to threshold.

### 3.4 Generality of the model

To test how the model generalizes, first we performed independent fits at each instant of the game, which correspond to different fractions of time available, and exhibit different RT distributions. Every distribution was fitted with the proposed model, obtaining *R*, *ρ* and Δ*T* parameters, and compared by their median value and a two-sample Kolmogorov-Smirnoff test of the complete distribution. In Fig. 2b, we show the median value obtained both from players and model data at each instant of the game. We observe an almost perfect match to the proposed model with obstacles. To quantify this observation, at each instant of game, we verified the goodness of fit comparing the distribution of human (black line) and simulated data (red line: no obstacles model; blue line: obstacle model) by analyzing JS divergence between distributions (see Fig. 2d). Results show that all RT distributions are indistinguishable from the best fit of model with obstacles (Fig. 2d, blue dots); a KS test between obstacles model and human data is non-significant in every instant of the game.

In contrast, when testing the DDM versus human data with the KS test, the null hypothesis is rejected (*p* < 10^−3^) indicating that the basic model without obstacles cannot reproduce human RT behavior. In accordance to this result, JS divergence show much higher values than obstacles model (Fig. 2d). We conclude that our obstacle model is universal to the decision-making process when playing on-line chess, regardless the instant of the game.

We further investigated whether this model extends to different cognitive domains. For this, we analyzed navigation logs of users in MercadoLibre^2^, the biggest on-line buying website of Latin-America. Logs consist of response times of users after doing a product search, i.e. how long users take to decide for (more information of) a product. Figure 4 shows the RT in this website; again, with a super-exponential tail. The obstacle model is plotted in blue line showing an almost perfect match (KS test, *D_n_* = 0.018). This result suggests that the model may transfer to very different domains (from chess to on-line buying), and that it is reflecting a intrinsic decision-making process.

**Fig 4.**
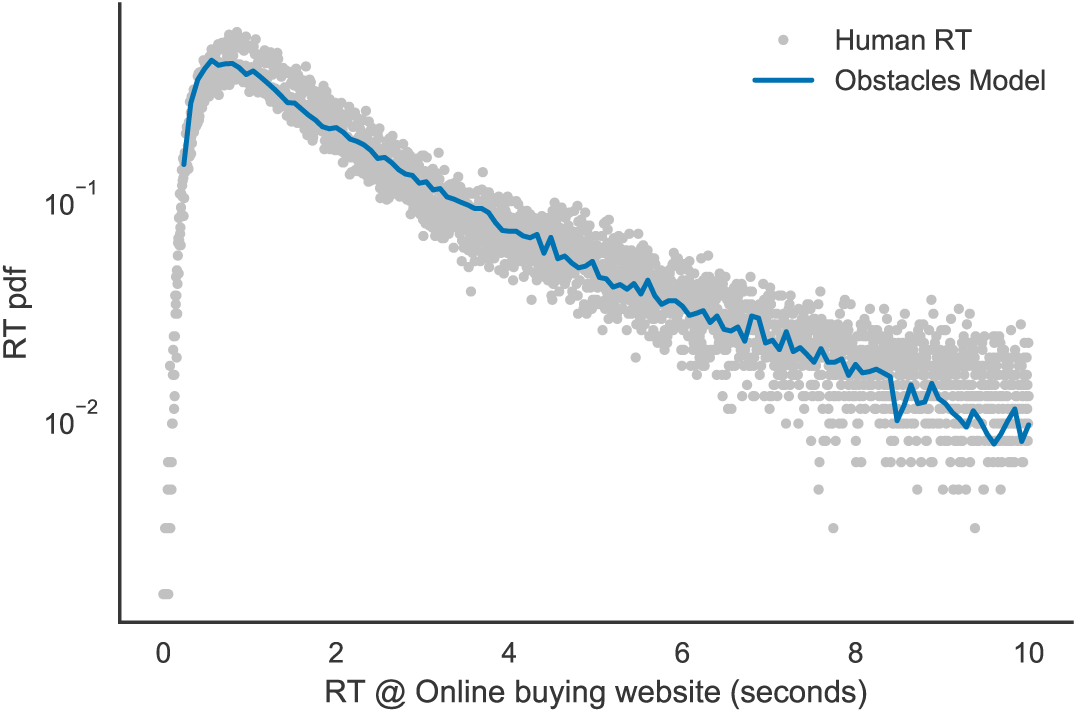
Generality. Model extends to on-line buying decision-making. Dots show response times between clicks of users doing a product search, i.e. how long users take to decide for (more information of) a products; blue line shows best RT fit of model showing an almost perfect match (Kolmogorov-Smirnoff Test, *D_n_* < 10^−8^, *p* > 0.99).

## 4 Discussion

Different models of accumulation to a bound have been the hallmark of two-choice decision-making [1, 34, 35]. A few studies have investigated experimentally (e.g [36, 37]) how they extend to more than two alternatives. For instance, Churchland and colleagues studied a four-choice paradigm [36]. Their results show that an urgency signal was more prominent on the two-choice paradigm than four-choice, consistent with longer response times on the latter experiment. Theoretical investigations have generalized competing integrators of higher number of alternatives [36, 38, 39]. Others have modeled by a competition between neural pools [5, 13]. These studies have succeeded in accounting for a range of behavioral data in conscious multi-choise tasks [40].

In chess, as in many other domains of human problem solving, it is well known that participants do not exhaustively search all alternatives [41]. Actually, studies show that boards evaluation is rather small, and only a few moves simulations are evaluated by players [42]. Chess players search in a decision tree halting this process by evaluations or heuristics which dictate that a given state is not desirable [43]. How decisions emerge without an exhaustive exploration of move alternatives is still an open question. Current chess models suggest that cognitive architecture should concentrate on relevant pieces or positions in the board, and may search for the best move by analogy with previous studied/plausible positions [44].

Classic DDM models are unable to capture the RT distributions produced by chess plays. Our model is designed to capture this process by presenting entropic barriers which can be seen as stop points in a search path, i.e. a moment in which there is sufficient visibility to discard the path based on an evaluation function. However, this interpretation must not be overstated. In a 2D grid the number of branches at each state is bound in 4 (all possible directions). This limit if obviously not true in chess or other multiple-choice tasks, but resembles the idea of having multiple trajectories available at each iteration in the decision process.

Using chess as a model of expertise in decision-making, Gobet and colleagues have demonstrated that stronger players do not necessarily outperform the maximum search depth of decision trees than weaker ones [45]. Instead, they cover deeper searches in average, showing a more exhaustive coverage of branches and a very efficient cutting threshold. In agreement with this view, we show that fitting RT distributions with our three-parameter *entropic* model achieves a remarkable precision. Our model refines this idea indicating that better players explore solutions in a more punctuated space. We interpret the model’s entropic barriers as stop points in a search process. The fact that stronger players show an increased number of obstacles indicates a more efficient cut algorithm for discarding sub-optimal branches. Expert players compensate the increase in time due to a navigation in a more punctuated space by a smaller distance to the threshold.

The model of decision-making we present here departs from traditional approaches by explicitly incorporating the presence of such entropic barriers in a stochastic search process. Using rapid chess big data as an unprecedented, high-throughput experimental laboratory, we show that our model provides a remarkable fit to the response time statistics (i.e. the distribution of times-to-move) at different stages of the game, not only first and second order moments but for the entire probability distribution over several response-time orders of magnitude. While at present we do not have the tools to investigate the potential mapping of the formal solutions to mental processes, it is expected that traces of them, in particular the presence of entropic barriers, should be found in the evidence that is readily available in psychophysical measurements.

## Acknowledgments

This research was supported by UBA, CONICET, ANPCyT and Human Frontiers Program. MS is sponsored by the James McDonnell Foundation. DFS is sponsored by Microsoft Faculty Fellow program. We thank the developers and administrators of FICS and FICSGAMES for continuously supporting a free chess server of the highest quality and completely open to experimentation and Carlos Diuk for fruitful discussions and useful comments on the manuscript.

## Footnotes

1 FICS: http://www.freechess.org/

2 http://www.mercadolibre.com.ar/

